# Characterization of SdGA, a cold-adapted glucoamylase from *Saccharophagus degradans*

**DOI:** 10.1101/2021.02.19.431967

**Authors:** Natael M. Wayllace, Nicolas Hedín, María V. Busi, Diego F. Gomez-Casati

## Abstract

We investigated the structural and functional properties of SdGA, a glucoamylase (GA) from *Saccharophagus degradans*, a marine bacterium which degrades different complex polysaccharides at high rate. SdGA is composed mainly by a N-terminal GH15_N domain linked to a C-terminal catalytic domain (CD) found in the GH15 family of glycosylhydrolases with an overall structure similar to other bacterial GAs. The protein was expressed in *Escherichia coli* cells, purified and its biochemical properties were investigated. Although SdGA has a maximum activity at 39°C and pH 6.0, it also shows high activity in a wide range, from low to mild temperatures, like cold-adapted enzymes. Furthermore, SdGA has a higher content of flexible residues and a larger CD due to various amino acid insertions compared to other thermostable GAs. We propose that this novel SdGA, is a cold-adapted enzyme that might be suitable for use in different industrial processes that require enzymes which act at low or medium temperatures.

## 1. Introduction

Glucoamylases (GAs) are hydrolytic enzymes also known as amyloglucosidases, glucan 1,4-alpha-glucosidases or exo-1,4-α-glucosidases, EC 3.2.1.3. GAs hydrolyze glycosidic α-1,4 bonds (but also α-1,6 bonds) from the non-reducing ends of starch molecules and maltooligosaccharides releasing β-D-glucose. These are typically microbial enzymes present in archaea, bacteria and fungi but absent in animals and plants, and they are classified into the GH15 family of glycoside hydrolases (www.cazy.org) [1, 2].

Glucoamylases from the GH15 family are multi-domain enzymes characterized by having a catalytic domain (CD) with a barrel-shaped (α/α)_6_, sometimes bound to a non-CD [1]. Depending on its origin, this non-CD could be a carbohydrate binding module (CBM) from CBM20 or CBM21 families (found, for example, in the GAs from *Aspergillus niger* or *Rhyzopus oryzae*, respectively) [3, 4]. However, there are other GA proteins that lack associated modules (such as that from *Sacharomycopsis fibuligera*) or have a particular N-terminal domain enriched in serine or threonine residues such as the GA from *Saccharomyces cerevisiae* [1]. Finally, most prokaryotic GAs contain the characteristic (α/α)_6_ barrel CD with a N-terminal β-sandwich domain such as the enzyme from *Thermoanaerobacterium thermosaccharolyticum* [5].

The main application of GAs (sometimes together with α-amylases and pullulanases) occurs in the process of saccharification of partially processed starch or dextrins to obtain glucose. Starch is the source of reserve carbohydrates in plant cells and algae and is one of the main renewable resources in nature [6–9] and it is composed by glucose molecules linked through a glycosidic bond. Starch consists mainly of two homopolysaccharides: amylopectin and amylose. Amylopectin is a polymer of D-glucose linked by α-1,4 bonds with about 5% branching with α-1,6 bonds, whereas amylose is essentially a linear polymer of glucose linked by α-1,4 glycosidic bonds with a small number of ramifications [7]. Starch is also a major component of most of the world’s staple foods such as rice, corn, potatoes, beans, wheat and cassava. In order to be used in the human diet, starch is frequently subjected to both chemical and enzymatic transformations to produce a wide variety of products, such as starch hydrolysates, glucose syrups, fructose, maltodextrin derivatives or cyclodextrins, which are used largely in the food industry [10]. In addition, starch is also widely used in other industries such as for the production of many textiles, pharmaceuticals, biodegradable plastic industry and in the production of biofuels, mainly bioethanol [11].

Over several years, many GA overproducing strains have been isolated in order to obtain high quantities of the enzyme. However, a more targeted strategy at the present is the cloning of specific genes or the discovery of new enzymes and their overexpression in heterologous organisms [10]. In the last decades, GAs from hyperthermophilous archaea of the genus *Sulfolobus*, *Thermoplasma*, *Picrophilus* and *Methanoccocus* have been investigated and characterized, and showed a very high thermostability and optimum temperature but a low specific activity [12–14]. In contrast, most of the GAs from fungi have optimal activity at 45°C and low pH [10, 15, 16]. Some fungal glucoamylases such as those from *Aspergillus* strains are widely used in industrial processes for starch hydrolysis because they display high activity at temperatures between 45 and 60°C [10].

Currently, there is strong interest in finding GAs with a better performance at low temperatures because these enzymes would avoid the heating requirement in some industrial processes such as starch saccharification among others, and, in this way, production costs could be minimized [17].

Recently, many amylases from marine bacteria have been characterized and proposed as a good alternative for starch degradation [18–21]. *Saccharophagus degradans* is a gram-negative bacterium that was isolated from a marsh grass, *Spartina alterniflora*, found at the bottom of the Chesapeake Bay, MD, USA [22]. *S. degradans* is able to degrade at least ten complex polymers such as agar, alginate, chitin, cellulose, fucoidan, laminarin, pectin, pullulan, starch and xylan [22]. These polymers are derived from numerous sources, such as algae, land plants, crustaceans, bacteria, and fungi. The high degradation rate of the different polysaccharides shown by *S. degradans* makes this bacterium a good candidate to obtain and characterize the properties of the enzymes that degrade these polymers.

In this way, we decided to carry out the structural and functional characterization of a putative GA from *S. degradans* (SdGA). The ability of SdGA to catalyze the reaction at low and medium temperatures, as well as its low thermal stability and increased amino acid flexibility compared to other more thermostable GAs, allow it to be classified as a cold-adapted enzyme. We propose that SdGA constitutes a good alternative for different industrial processes such as starch saccharification at low or mild temperatures and its subsequent application in biofuel production, mainly bioethanol.

## 2. Materials and Methods

### 2.1 Construction of the recombinant expression vector and transformation of *E. coli* cells

The cDNA sequence coding for the mature form of the GA from *S. degradans* (ABD79864.1), named *SdGA,* was synthesized and cloned into the pRSF-DUET vector by Genscript Biotech Corp. (Piscataway, NJ, USA) and the construction was named pRSF-DUET_SdGA. *SdGA* was synthesized with codon optimization for heterologous expression in *E. coli* cells (GenBank MT754655) and was fused to a 5‘ sequence coding for a His_6_ tag at the N-terminal region of the SdGA protein.

### 2.2 Expression and purification of SdGA

*E. coli* BL21(DE3) Rosetta was transformed with pRSF-DUET_SdGA and colonies were selected in LB agar plates containing 25 mg/ml of kanamycin and 50 mg/ml of chloramphenicol. Transformed cells were cultured in LB media containing the same antibiotics at 37°C to *A*_600_ ~ 0.5-0.6. Protein expression was induced at 37°C with isopropyl-β-D-thiogalactopyranoside (IPTG) at a final concentration of 1 mM for 2 h. The cells were harvested by centrifugation at 5,000 rpm for 10 min., resuspended in 20 mM sodium phosphate buffer (pH 7.0) at 4°C and then disrupted with an ultrasonicator (VCX130, Sonics and Materials Inc., Newtown, CT, USA). The homogenate was centrifuged at 5,000 rpm for 20 min. Recombinant His-tagged-SdGA was purified from the soluble fraction using a HiTrap Chelating column (HiTrapTM Chelating HP, Amersham Biosciences) equilibrated with 0.1 M NiSO_4_. The protein was eluted using a discontinuous gradient of increasing imidazole concentration (20 to 500 mM). The purified enzyme was concentrated and stored at −20°C and was found to be stable for at least 2 months.

### 2.3 Sequence analysis, flexibility evaluation and homology modeling of SdGA

Similarity search of SdGA was performed using the BLASTp algorithm at the National Center for Biotechnology Information (NCBI) database (https://blast.ncbi.nlm.nih.gov/Blast.cgi). Computer-assisted protein sequence analyses were performed using ClustalW version 2.0. [23]. The phylogenetic tree for SdGA was constructed using maximum likelihood method based on the JTT matrix-based model with Bootstrap method phylogeny test and 500 replications using MEGA7 program [24, 25].

The amino acid sequence of SdGA and other GAs were analyzed and compared using PSSM-based encoding method from FlexPred server (http://flexpred.rit.albany.edu) to evaluate the flexibility or rigidity of the residues in the CD [26]. The percentage of glycine (%G) and flexible residues (%F) within the CD were calculated.

3D structural model was obtained using the Modeller @TOME V3 program [27] using T.I.T.O. (Tool for Incremental Threading Optimization) for the alignment between query and template [27]. The 3D model of SdGA was built using the structure of a GA from *Thermoanaerobacterium thermosaccharolyticum* as a template (PDB entry: 1LF6, 34% identity). The quality of the model was evaluated with ProSA-web structure analysis program [28, 29] and Verify 3D [30]. Superposition of SdGA/tGA was performed using PyMOL software (The PyMOL Molecular Graphics System, Version 2.0 Schrödinger, LLC, https://pymol.org/2/).

### 2.4 Glucoamylase activity assay and kinetic analysis

Glucoamylase activity was measured by determining free glucose released after maltose hydrolysis. The reaction mixture consisted of 0.1 M sodium phosphate buffer (pH 6.0) containing 2% (w/v) maltose, 0.032 μg/μl of SdGA at final volume 100 μl at 39°C for 20 min. The amount of released glucose was determined using a commercial kit based on the method of glucose oxidase (GOD)/peroxidase (POD) (Wiener laboratories, Rosario, SF, Argentina) [31]. One unit of GA hydrolytic activity was defined as the amount of enzyme needed to release 1 μmol of glucose per min. at 39°C. Assays were performed at least by triplicate at different enzyme concentrations to ensure steady-state conditions. Protein concentrations were estimated according to the method of Bradford [32].

The effect of maltose concentration on SdGA activity was evaluated in 0.1 M sodium phosphate buffer (pH 6.0) at 39°C. The kinetic parameters *K_m_*, *k*_cat_, *S*_0.5_, *V*_max_ and *n*_H_ were calculated with a computer program using the Levenberg-Marquardt algorithm for regression by fitting the data to the Hill equation [33]. *S*_0.5_ is defined as the concentrations of substrate that give 50% maximal activity (*V*_max_), *k*_cat_ is the turnover number and *n*_H_ is the Hill number.

The effect of pH on the activity of SdGA was determined in two 0.1 M buffers with maltose as the enzyme substrate (sodium acetate buffer, pH 4 - 5.6 and sodium phosphate buffer, pH 6 - 8.0, respectively) at 39°C. The effect of temperature on SdGA activity was determined with 2% (w/v) maltose between 10– 50°C as described above. Temperature stability assays were performed by incubating the same quantity of SdGA at different temperatures for 0 to 30 min., and the residual activity was measured under standard assay conditions. The effect of metabolites on the thermal stability of SdGA was measured using 0.1 mM acarbose (Sigma-Aldrich cat. n° 56180-94-0), 1 mM CaCl_2_ or 10% (v/v) glycerol. The first order equation ln (A/A_0_) = −*k*. t was used to calculate the inactivation rate constant (*k*, min^−1^). A is the residual activity at time t and A_0_ is the initial activity at time zero. *D-*value (Decimal reduction time), the time to reduce the initial activity 90%, was calculated from the equation: *D* = ln(10) / *k* as described [34]. All measurements were made at least by triplicate. Significant differences were determined by one-way ANOVA and Dunnet test using GraphPad Prism version 5.0 software (GraphPad Software, La Jolla, CA, USA)

### 2.5 Additional methods

The purified SdGA was analyzed by SDS-PAGE using 12% (w/v) gels as described by Laemmli [35] with a Bio-Rad Mini Protean system (Bio-Rad, Hercules CA, USA). Gels were stained by Coomassie Blue and/or electro-blotted onto nitrocellulose membranes (HybondTM-ECLTM, Amersham Biosciences, UK). Electroblotted membranes were incubated with penta-His antibody (Qiagen, Valencia, CA, USA). The antigen–antibody complex was visualized with alkaline phosphatase-linked to anti-rabbit IgG followed by staining with 5-bromo-4-chloroindol-2-yl phosphate and nitro blue tetrazolium [36].

## 3. Results

### 3.1 Sequence analysis and domain identification

The analysis of the nucleotide sequence of SdGA shows the presence of an open reading frame of 2,409 bp coding for a putative GH15 GA. The protein is composed by 803 amino acids with a calculated molecular mass of 87.4 kDa. The analysis using SignalP-5.0 Server (http://www.cbs.dtu.dk/services/SignalP/) revealed the presence of a 20 amino acids signal peptide (SP) in the N-terminus of SdGA. Using InterPro v.79.0 server (http://www.ebi.ac.uk/interpro/) we found that SdGA contains a family 15 glycoside hydrolase N-terminal domain (GH15_N, amino acids 38-322), uniquely found in bacterial and archaeal glucoamylases and glucodextranases, and a C-terminal GH15-like CD (GH15L-CD, amino acids 340-781) similar to the six-hairpin domain present also in the GH15 family (Figure 1a). The CD contains six α-hairpins arranged in a closed circular array named α/α toroid fold, [37]. Both domains are connected by a linker region (LR) composed by two α-helices, according to the analysis with Jpred 4 (http://www.compbio.dundee.ac.uk/jpred/) [38].

**Figure 1.**
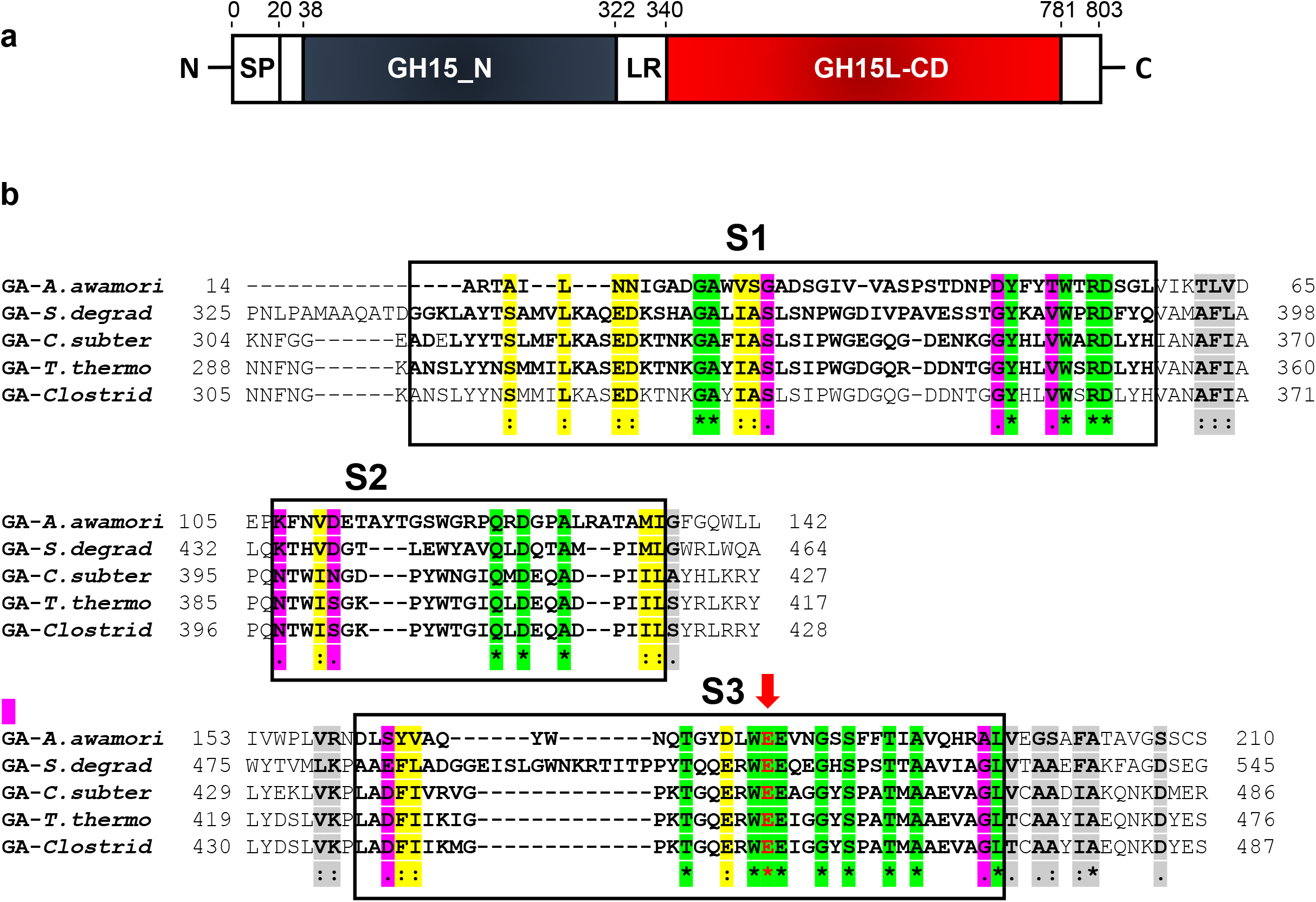
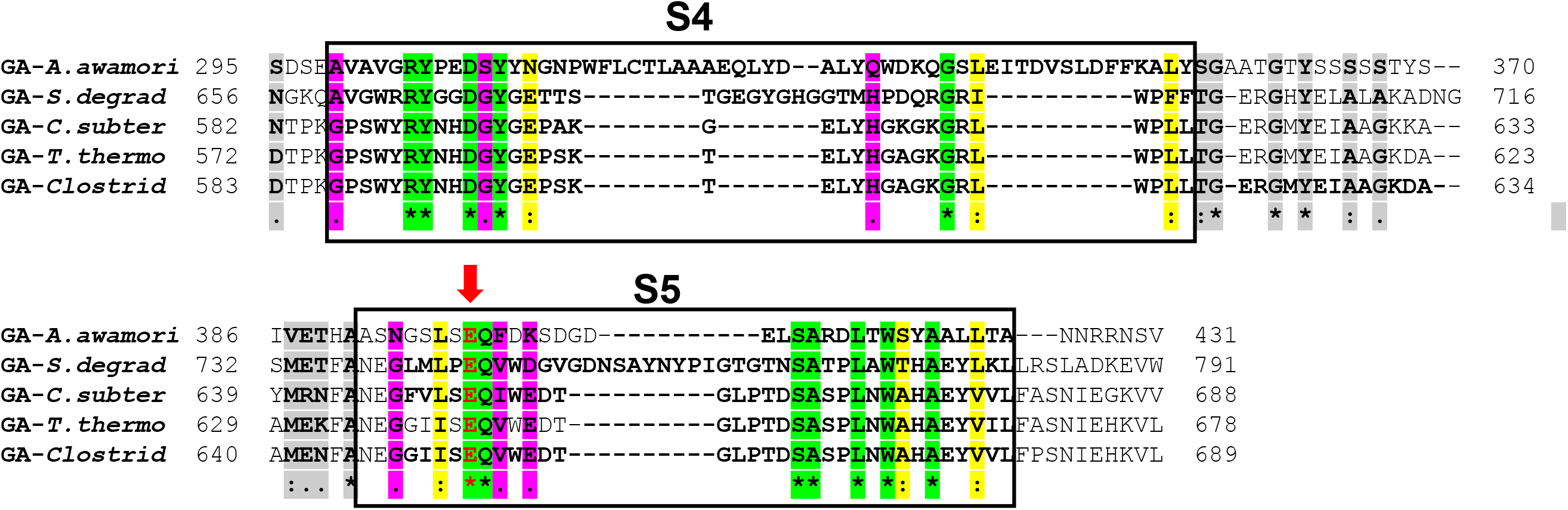
Structure domain and sequence alignment of SdGA. a) Structure domain of SdGA: SP: signal peptide, GH15_N: glycoside hydrolase family 15 N-terminal domain, LR: linker region, GH15L-CD: C-terminal glycoside hydrolase family 15-like CD. Numbers at the top indicate the amino acid positions at the beginning and end of each domain b) Sequence alignment of the CD of SdGA with other GAs: *Aspergillus awamori* (PDB entry: 1GLM); *Saccharophagus degradans* (SdGA) ABD79864.1; *Caldanaerobacter subterraneus* subsp. *tengcongensis* AAM25005.1; *Thermoanaerobacterium thermosaccharolyticum* (PDB entry: 1LF6) and *Clostridium sp.* BAA02251.1. Boxes show the five highly conserved regions (S1–S5) found in GAs. Red arrows indicate two catalytic residues, E513 and E745 (SdGA numbering). The conserved residues are marked with * (green inside the rectangles or gray outside them) and with: and . (pink and yellow) the conservative substitutions.

The amino acid sequence of SdGA displayed the highest sequence identity with marine GAs from *Gilvimarinus polysaccharolyticus* (WP_049722861.1, 67% identity), and *Gilvimarinus agarilyticus* (WP_041522105.1, 66% identity). Figure 1b showed a multiple sequence alignment of the CDs of SdGA and several other known bacterial and fungal GAs. We found that SdGA contained five conserved regions (S1 to S5, boxed in Figure 1b) typically found in the CD of GAs defined previously [39]. In addition, SdGA contains two conserved glutamate residues, E513 and E745 (marked with red arrows, Figure 1b), that are the putative catalytic acid and base, which correspond to the E438 and E636 residues in *T. thermosaccharolyticum* GA [5].

Based on the amino acid sequence of SdGA, we have constructed a phylogenetic tree using the maximum likelihood method. Clearly, SdGA showed phylogenetic similarity with other GAs from marine bacteria but switched to different clusters away from other non-marine bacterial species, such as *Clostridium* sp. and also fungi and archaea, indicating the divergence of GAs among organisms (Figure 2).

**Figure 2.**
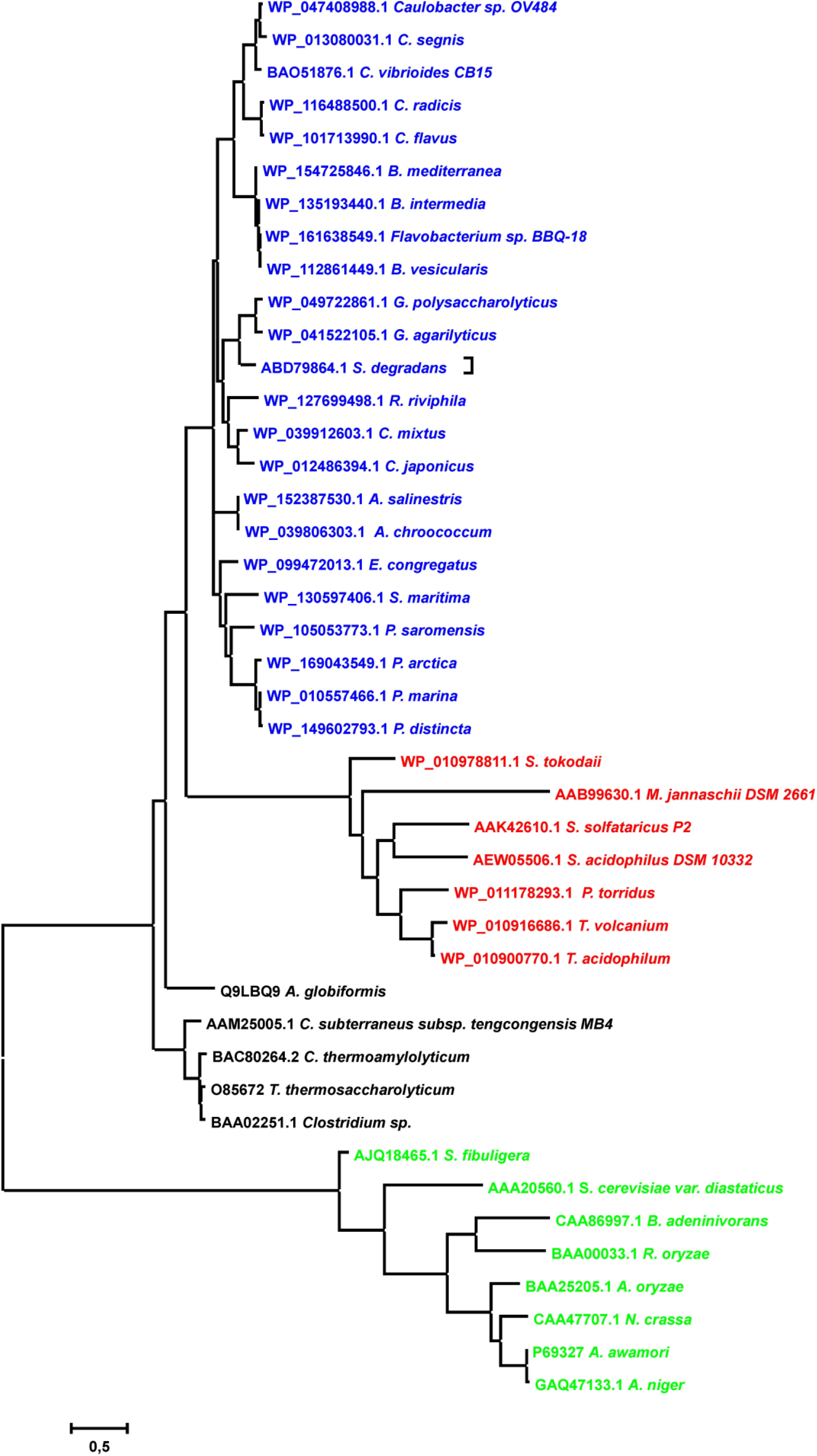
Phylogenetic Analysis of GAs. Phylogenetic tree showing the relationship among GAs from different microorganisms based on amino acid sequence homology. Multiple sequence alignments were done using Clustal W program and the phylogenetic analysis was carried out by Maximum Likelihood method. The unrooted phylogenetic tree is shown using MEGA7 program [24]. The sequences were retrieved from NCBI (http://www.ncbi.nlm.nih.gov/). The accession numbers are: WP_047408988.1 glucan 1,4-alpha-glucosidase *Caulobacter sp*. *OV484*, WP_013080031.1 glucan 1,4-alpha-glucosidase *Caulobacter segnis*, BAO51876.1 glucoamylase *Caulobacter vibrioides* CB15, WP_116488500.1 glucan 1,4-alpha-glucosidase *Caulobacter radicis*, WP_101713990.1 glucan 1,4-alpha-glucosidase *Caulobacter flavus*, WP_154725846.1 glucan 1,4-alpha-glucosidase *Brevundimonas mediterranea*, WP_135193440.1 glucan 1,4-alpha-glucosidase *Brevundimonas intermedia*, WP_161638549.1 glucan 1,4-alpha-glucosidase *Flavobacterium sp. BBQ-18*, WP_112861449.1 glucan 1,4-alpha-glucosidase *Brevundimonas vesicularis*, WP_049722861.1 glucan 1,4-alpha-glucosidase *Gilvimarinus polysaccharolyticus*, WP_041522105.1 glucan 1,4-alpha-glucosidase *Gilvimarinus agarilyticus*, ABD79864.1 glucoamylase *Saccharophagus degradans* (marked), WP_127699498.1 glucan 1,4-alpha-glucosidase *Rheinheimera riviphila*, WP_039912603.1 glucan 1,4-alpha-glucosidase *Cellvibrio mixtus*, WP_012486394.1 glucan 1,4-alpha-glucosidase *Cellvibrio japonicus*, WP_152387530.1 glucan 1,4-alpha-glucosidase *Azotobacter salinestris*, WP_039806303.1 glucan 1,4-alpha-glucosidase *Azotobacter chroococcum*, WP_099472013.1 glucan 1,4-alpha-glucosidase *Emcibacter congregatus*, WP_130597406.1 glucan 1,4-alpha-glucosidase *Shewanella maritima*, WP_105053773.1 glucan 1,4-alpha-glucosidase *Psychrosphaera saromensis*, WP_169043549.1 glucan 1,4-alpha-glucosidase *Pseudoalteromonas arctica*, WP_010557466.1 glucan 1,4-alpha-glucosidase *Pseudoalteromonas marina*, WP_149602793.1 glucan 1,4-alpha-glucosidase *Pseudoalteromonas distincta*, WP_010978811.1 glucoamylase *Sulfurisphaera tokodaii*, AAB99630.1 glucoamylase *Methanocaldococcus jannaschii DSM 2661*, AAK42610.1 glucoamylase *Sulfolobus solfataricus* P2, AEW05506.1 glucoamylase *Sulfobacillus acidophilus DSM 10332*, WP_011178293.1 glucoamylase *Picrophilus torridus*, WP_010916686.1 glucoamylase *Thermoplasma volcanium*, WP_010900770.1 glucoamylase *Thermoplasma acidophilum*, Q9LBQ9 glucodextranase *Arthrobacter globiformis*, AAM25005.1 glucoamylase *Caldanaerobacter subterraneus subsp. tengcongensis MB4*, O85672 glucoamylase *Thermoanaerobacterium thermosaccharolyticum*, BAC80264.2 glucoamylase *Clostridium thermoamylolyticum* BAA02251.1 glucoamylase *Clostridium sp.*, AJQ18465.1 glucoamylase *Saccharomycopsis fibuligera* AAA20560.1, glucoamylase *Saccharomyces cerevisiae var. diastaticus*, CAA86997.1 glucoamylase *Blastobotrys adeninivorans*, BAA00033.1 glucoamylase *Rhizopus oryzae*, BAA25205.1 glucoamylase *Aspergillus oryzae*, CAA47707.1 glucan 1,4-alpha-glucosidase *Neurospora crassa*, P69327 glucoamylase *Aspergillus awamori* and GAQ47133.1 glucoamylase *Aspergillus niger*. Blue: marine bacterial GAs, black: non-marine bacterial GAs, red: archaeal GAs, green: fungal GAs

### 3.2 Homology modelling of SdGA

A homology model of SdGA protein was built as described under Materials and Methods section using the 3D structure of tGA, a GA from *Thermoanaerobacterium thermosaccharolyticum* (PDB entry: 1LF6, 34% identity). Analysis using Verify 3D program showed that 81.6% of the residues have averaged 3D-1D score higher than 0.2, whereas a Z-score of −7.05 was obtained using ProSA-web. According to these results, we found that the SdGA model has a good quality.

The SdGA model exhibited a fold similar to tGA with both α-helical and β-sheet secondary structures conserved (Figure 3a). Three structurally different regions can be distinguished, (i) the N-terminal domain, which is made up of 16 antiparallel β-strand segments divided into 2 β-sheets; (ii) the 2 α-helice linker region and (iii) the C-terminal domain, which has, as described above, an (α/α)_6_ barrel structure similar to other GAs (Figure 3a).

**Figure 3.**
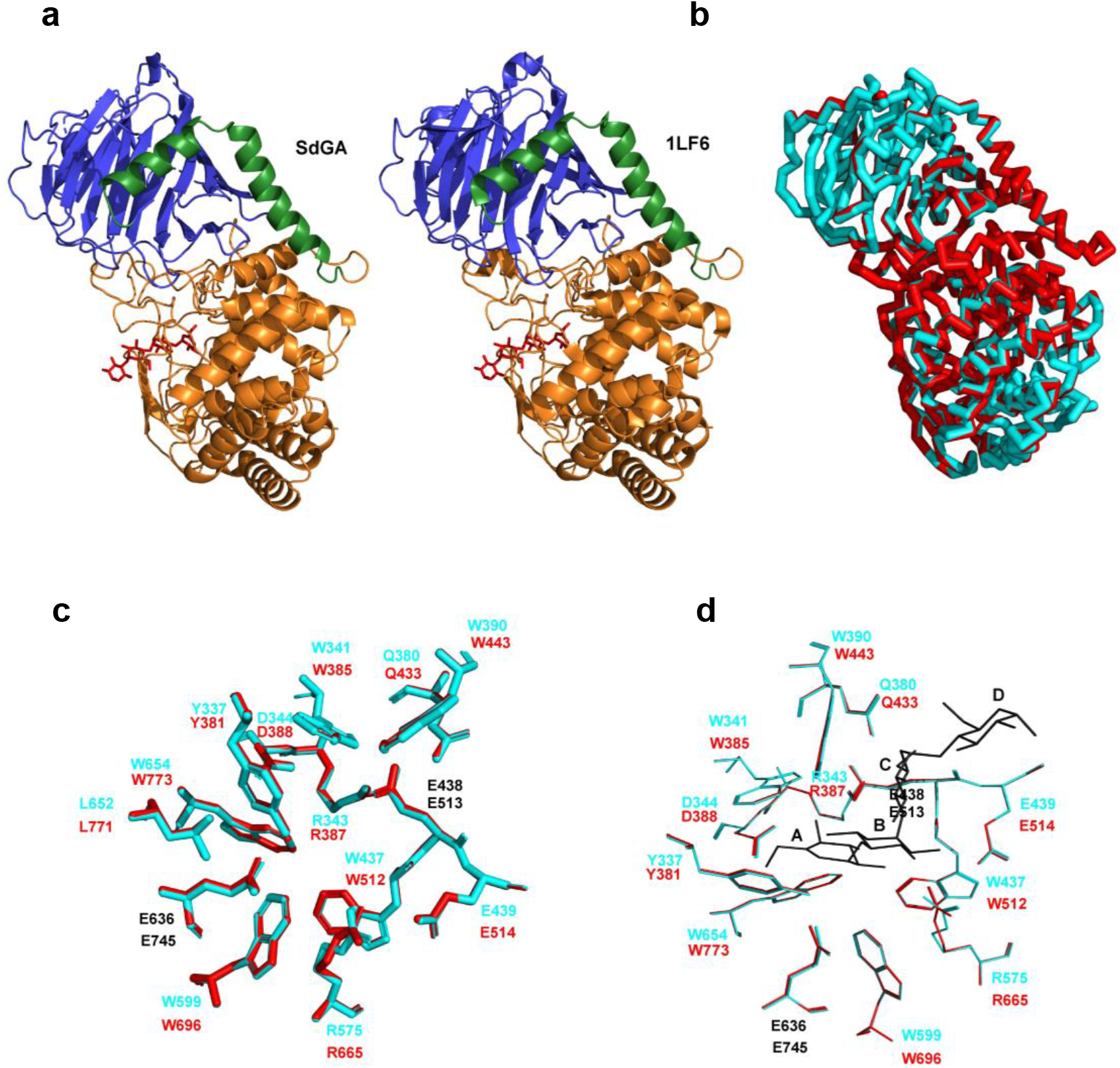
Homology Modelling of SdGA. a) Proposed model for SdGA (left) and structural model of *T. thermosaccharolyticum* GA (PDB entry: 1LF6, right). b) Superposition of SdGA (red) and 1LF6 (cyan). Superposition between SdGA model and 1LF6 structure showing the residues involved in the active site in absence (c) and presence of acarbose (d). SdGA residues are shown in red, 1LF6 residues are shown in cyan and the catalytic residues are marked in black. In d, bound acarbose is shown in black and A, B, C and D on the acarbose molecule represents the four subunits of the tetrasaccharide.

The superimposition of the polypeptide backbone structures of SdGA and tGA showed that both 3D structures are also very similar (Figure 3b). It was determined that the amino acids postulated by Aleshin and colleagues [5] as belonging to the active site and acarbose binding (Y337, W341, R343, D344, Q380, W390, W437, E438, E439, W599, R575, E636, W654 and L652) are well conserved in the SdGA model (Figure 3c, 3d and Table 1).

**Table 1.** Amino acid residues involved in catalysis and acarbose binding in tGA and SdGA.

| tGA | SdGA | Function |
| --- | --- | --- |
| Y337 | Y381 | Close contact (3.3 Å) with C7. Unit A of acarbose |
| W341 | W385 | Close contact (3.5 Å) with C6. Unit A of acarbose |
| R343 | R387 | H-bond O3 and O4. Unit A of acarbose |
| D344 | D388 | H-bond O6. Unit A of acarbose |
| Q380 | Q433 | H-bond O6. Unit C of acarbose |
| W390 | W443 | Hydrophobic interactions with Unit C and D of acarbose |
| W437 | W512 | H-bond O3. Unit B of acarbose |
| E438 | E513 | Catalytic acid residue. |
| E439 | E514 | H-bond O2. Unit B of acarbose |
| R575 | R665 | H-bond O2. Unit A of acarbose/H-bond O3. Unit B of acarbose |
| W599 | W696 | Contribute to the formation of a large hydrophobic wall |
| E636 | E745 | Catalytic base residue |
| W654 | W773 | Contribute to the formation of a large hydrophobic |

Ref:, O2, O3 and O6 oxygen atoms in acarbose, C6 and C7 carbon atoms in acarbose, Unit A, Unit B and Unit C are subunits of acarbose. H-bond means that an hydrogen bond is stabilized. tGA: GA from *Thermoanaerobacterium thermosaccharolyticum*.

### 3.3 Expression and purification of SdGA

SdGA from *S. degradans* was synthesized and cloned into the pRSFDuet vector, as described in the Materials and Methods section. The plasmid named pRSF-DUET_SdGA was successfully expressed in *E. coli* BL21(DE3) Rosetta cells. The expression was optimized to obtain about 40% of SdGA in the soluble fraction (Figure 4, lane 5). Thus, soluble SdGA was purified to homogeneity by a single purification step using Ni^2+^ affinity chromatography. Using this procedure, we have obtained about 0.2 mg of protein from about 5 g of *E. coli* cells. SdGA has a calculated molecular mass of 87.4 kDa and produced a single band of about 90kDa on SDS-PAGE analysis (lane 8, Figure 4). The identity of the protein band was confirmed by immunoblotting using anti-penta-His antibodies (lane 9, Figure 4).

**Figure 4.**
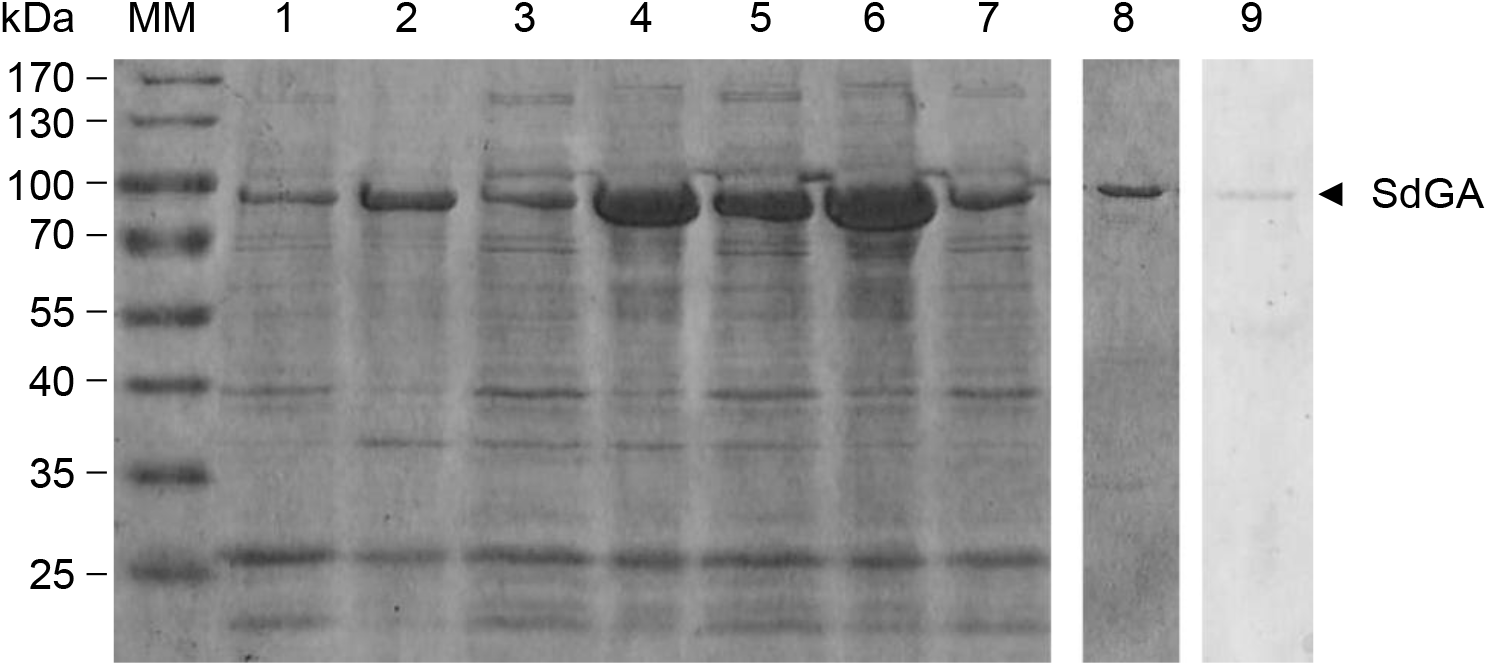
Expression analysis and purification of recombinant SdGA from *E. coli* cells. SDS-PAGE analysis followed by Coomasie Blue stain of: Lane 1: supernatant of crude extract from BL21 (DE3) Rosetta without induction; Lanes 2, 4 and 6: insoluble proteins after 1, 2 and 3 h of induction, respectively; Lanes 3, 5 and 7: soluble proteins after 1, 2 and 3 h induction, respectively; Lane 8: recombinant purified SdGA. Lane 9: western blot analysis of SdGA followed by incubation with anti-penta-His antibodies. Numerals on the left indicate molecular masses (MM, in kDa) of the Pre-stained SDS-PAGE standard low range (Thermo Fisher Scientific).

### 3.4 Kinetic characterization of purified SdGA

Purified SdGA showed more than 80% glucoamylase activity at pH values between 5.5 and 6.5, being pH 6 the optimum for the enzyme. Moreover, SdGA remains active over the range between pH 4.0 and pH 8.0 (Figure 5a).

**Figure 5.**
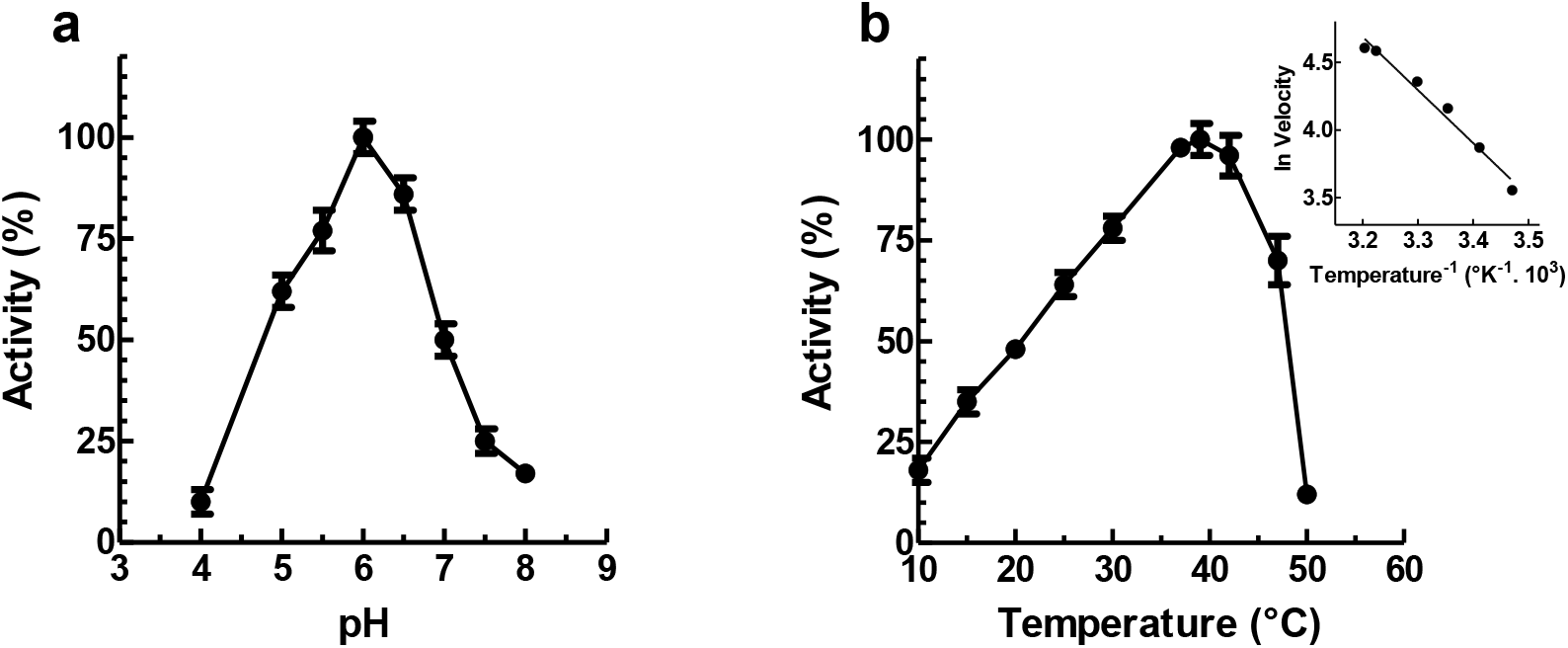
Effect of pH and temperature on the activity of SdGA. a) Effect of pH on the activity of recombinant SdGA. Activity was assayed with maltose as substrate using 0.1 M sodium acetate buffer (pH 3.6 – 5.5) or 0.1 M sodium phosphate buffer (pH 5.8 – 8.0), at 39°C. b) Effect of the temperature on the activity of SdGA. Activity was assayed with maltose as substrate using 0.1 M sodium phosphate buffer at pH 6.0. Inset: Arrhenius plots for the activity of SdGA assayed at different temperatures. All data are the means of 3 independent experiments ± SD.

The dependence of SdGA activity on the temperature was examined using 2% (w/v) maltose in 0.1 M sodium phosphate buffer (pH 6.0) at temperatures ranging from 10 to 50°C. Figure 5b shows that the activity of the SdGA was maximal at 39°C. However, we found that the enzyme catalyzes the reaction with at least 50% of the *V*_max_ between 20 to 48°C, showing about 20-30% of activity at 10-15°C. The enthalpy of activation for the reaction calculated from the Arrhenius equation is 31.8 kJ/mol (inset Figure 5b). Thus, SdGA is able to catalyze the reaction in a wide range of temperatures as reported for other GAs, classified as cold-adapted enzymes, like a fungal GA from *Tetracladium* sp. [40] or the GA from *Caulobacter crescentus* [41].

To determine the kinetic parameters *S*_0.5_, *n*_H_ and *V_max_* (or *k*_cat_, to compare with other GAs, see Table S1), GA activity was measured in 0.1 M sodium phosphate buffer (pH 6.0) at 20°C and 39°C, at increasing concentrations of maltose. Interestingly, SdGA shows no Ca^2+^ dependency for activity (not shown). At 39°C, SdGA showed Michaelis kinetics with a *V*_max_ = 6.1 ± 0.5 U/mg (*k*_cat_ = 8.9 ± 1.1 s^−1^), a *n*_H_ = 1.1 ± 0.1 and a *S*_0.5_ = 10.7 ± 1.5 mM., while at 20°C the enzyme showed a *V*_max_ and *S*_0.5_ about 2- and 4-fold lower (2.7 ± 0.22 U/mg and 2.5 ± 0.63 mM, respectively), resulting in a 2-fold increase of the catalytic efficiency respect to the condition at 39°C (*V*_max_/*S*_0.5_ = 0.56 and 1.1 U.mg^−1^. mM^−1^, respectively) (Figure 6).

**Figure 6.**
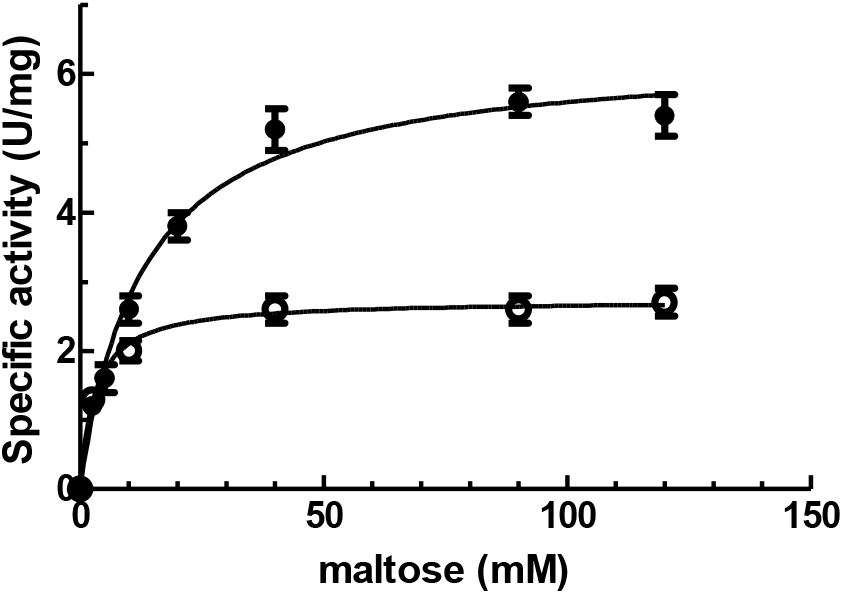
Maltose saturation plots for SdGA. SdGA activity was measured with increasing concentrations of maltose at 20°C (empty circles) or 39°C (black circles). See materials and methods for details.

### 3.5 Studies on thermal stability of SdGA

Temperature stability assays were performed by incubating different aliquots of the enzyme without the substrate at 45°C and 50°C at different times, and the residual activity was measured under standard assay conditions. As suggested by the results showed in Figure 7a, SdGA is unstable when exposed at temperatures above 45°C in the absence of effectors. Inactivation is dependent on temperature and incubation time and follows first-order kinetics. Inactivation half times (*t*_0.5_) obtained were 9.8 ± 0.7 and 0.8 ± 0.1 min for 45°C and 50°C, respectively (Figure 7a). The inactivation rate constants (*k*) at 45°C and 50°C were calculated from the slope of the inset of Figure 7a (0.071 ± 0.008 and 0.607 ± 0.0.83 min^−1^, respectively).

**Figure 7.**
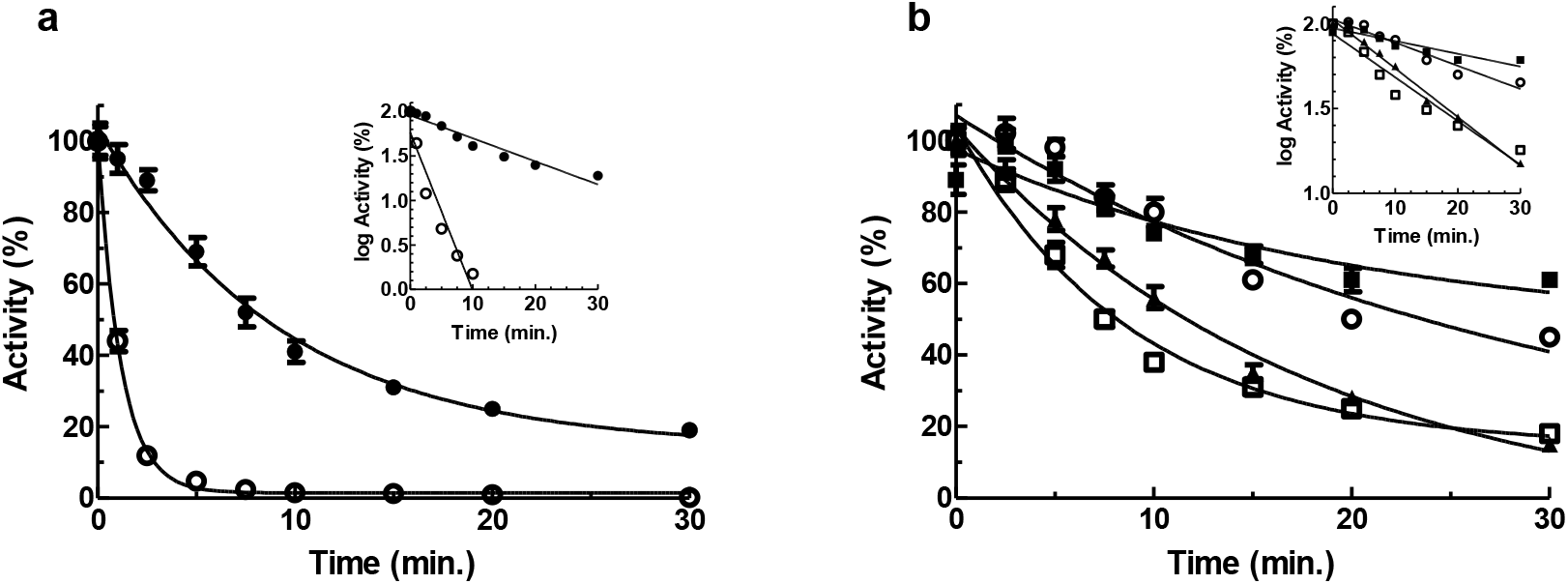
Thermostability of SdGA. (a) The enzyme was incubated at 45°C (black circles) or 50°C (empty circles) for different time intervals and the residual activity was measured at 39°C as described in Materials and Methods. The inactivation rate constants (*k*) were determined from the slopes of the logarithmic plot of activity vs. time (see inset). (b) Analysis of the thermostability of SdGA at 45°C in the absence (empty squares) or presence of different metabolites: 1 mM Ca^2+^ (black triangles), 10% (v/v) glycerol (empty circles) and 0.1 mM acarbose (black squares). The inactivation rate constants (*k*) for each condition were calculated as described in Fig. 7A (inset). All data are the means of 3 independent experiments ± SD.

We also tested the effect of different compounds on the thermal inactivation of SdGA at 45°C. Thus, we incubated the enzyme in the presence of 0.1 mM acarbose, 1 mM Ca^2+^ or 10% (v/v) glycerol at different times, and the remaining SdGA activity was measured at 39°C (Figure 7b). The semi-log plots of the residual activity vs. time were linear at all temperatures, (inset Figure 7b) indicating that the inactivation follows first-order kinetics for all the metabolites tested.

We found that acarbose shows partial protection as indicated by the increase of the time required to obtain 50% of the initial activity (*t*_0.5_ = 44.3 ± 2.5 min, while glycerol was less effective, showing a *t*_0.5_ of 21.2 ± 2.8 min. In contrast, Ca^2+^ does not significantly modified the *t*_0.5_ for the thermal inactivation of SdGA (*t*_0.5_ = 10.3 ± 1.9 min.), compared to the *t*_0.5_ showed by the enzyme incubated in the absence of effectors (9.8 ± 0.7 min.). The inactivation rate constants *k*, *D*-values and *t*_0.5_ are shown in Table 2. These data indicate that acarbose and to a lesser extent glycerol, are good protectors, albeit partially, against the thermal inactivation of the protein, while Ca^2+^ is a poor protective agent.

**Table 2.** *k*, *D* and *t*_0.5_ values for the thermal inactivation of SdGA in the presence of different compounds at 45°C.

| Compound | *k* (min<sup>-1</sup>) | *D* (min) | *t*<sub>0.5</sub> (min) |
| --- | --- | --- | --- |
| n.a. | 0.071 ± 0.008 | 32.4 ± 4.2 | 9.8 ± 0.7 |
| Ca<sup>2+</sup> 1 mM | 0.064 ± 0.004 | 35.9 ± 3.9 | 10.3 ± 1.9 |
| Glycerol 10% (v/v) | 0.029 ± 0.003 | 79.4 ± 6.8 | 21.2 ± 2.8 |
| acarbose 0.1 mM | 0.016 ± 0.002 | 143.9 ± 12.3 | 44.3 ± 3.5 |

Ref: *k*, thermal inactivation constant; *D*, Decimal reduction time; *t*<sub>0.5</sub>, half time of inactivation. Mean values ± SD are reported.

Results showed that SdGA is much thermolabile when compared with other characterized GAs. This characteristic, as well as the fact that the enzyme is active at low temperatures and that the activity rapidly decreases at temperatures over 42°C, allow this enzyme to be classified as a possible cold-adapted enzyme [40, 42, 43].

### 3.6 Analysis of flexible residues in SdGA

One of the properties of cold-adapted enzymes is their increased flexibility either in the vicinity of the active site or in more distant parts of the protein molecule. Moreover, this flexibility has been associated to a decrease in the structural thermostability of cold-active enzymes, which is in agreement with the thermal inactivation of SdGA under moderate temperatures reported above [42–45]. In addition, cold-adapted enzymes usually contain a higher number of glycines, mainly in loop regions, and a lower number of prolines and arginines [45].

Thus, we carried out a study in order to compare the flexibility and composition of amino acids present in SdGA with those of other GAs from psychrophilic, cold-adapted and thermophilic organisms. FlexPred analysis shows that the CD of SdGA has similar number of flexible residues (13.45 %F) compared with GAs from *B. mediterranea* (BmGA), *Caulobacter* sp. (CsGA) (about 15 %F) and *Pseudoalteromonas* species (about 14 %F, Table 3). This percentage is significantly higher than the percentage of flexible residues found in GAs from thermophilic bacteria such as those from *S. Solfataricus* (7.44 %F), *T. thermosaccharolyticum* (8.81 %F) or *T. acidophilum* (5.38 %F). In addition, SdGA, BmGA, CsGA and *Pseudoalteromonas* species have higher glycine content (between 10.5 – 11.7 %G) compared with the thermophilic GAs (between 5.5 – 9.7 %G). It is important to highlight that the GAs from *B. mediterranea*, *Caulobacter* sp., *Pseudoalteromonas* species and *Flavobacterium* sp. are grouped with SdGA within the same region of the phylogenetic tree shown in Figure 2.

**Table 3.** Percentage of flexible and glycine residues in the catalytic domain of various GAs.

| Enzyme (organism) | %G | %F |
| --- | --- | --- |
| SdGA | 10.51 | 13.45 |
| *Brevundimonas mediterranea* | 11.47 | 15.21 |
| *Caulobacter* sp. | 11.66 | 15.13 |
| *Pseudoalteromonas arctica* | 10.67 | 14.66 |
| *Pseudoalteromonas distincta* | 10.81 | 14.74 |
| *Pseudoalteromonas marina* | 10.76 | 14.42 |
| *Flavobacterium* sp. | 11.54 | 13.96 |
| *Sulfolobus solfataricus* | 5.47 | 7.44 |
| *Thermoplasma acidophilum* | 5.67 | 5.38 |
| *Thermoanaerobacterium thermosaccharolyticum* | 7.90 | 8.81 |
| *Caldanaerobacter subterraneus* subsp. *tengcongensis* | 9.73 | 8.58 |

Ref: %G and %F are the percentage of glycine and flexible residues, respectively, in the CD of each GA. Flexible residues were calculated using FlexPred (http://flexpred.rit.albany.edu). The accession number of GAs are: ABD79864.1 *Saccharophagus degradans*, WP\_154725846 *Brevundimonas mediterranea*, WP\_047408988.1 *Caulobacter* sp., WP\_169043549.1 *Pseudoalteromonas arctica*, WP\_149602793.1 *Pseudoalteromonas distincta*, WP\_010557466.1 *Pseudoalteromonas marina*, WP\_161638549.1 *Flavobacterium* sp. *BBQ-18*, AAK42610.1 *Sulfolobus solfataricus*, WP\_010900770.1 *Thermoplasma acidophilum*, AAC24003.1 *Thermoanaerobacterium thermosaccharolyticum*, AAM25005.1 *Caldanaerobacter subterraneus* subsp. *tengcongensis*.

It was suggested that other properties of cold-adapted enzymes are (i) their higher amino acid content compared to the enzymes of thermophilic organisms, which have a more compact core and (ii) the existence of amino acid inserts or loops in regions near the active site [41–43]. The latter would also be related to the thermolabile characteristics of this group of enzymes.

Using Clustal W, we performed a sequence alignment of the catalytic domains of SdGA with the GAs used to analyze the flexible residues. Results show the presence of 5 insertions between 9 to 13 amino acids that are absent in all analyzed thermophilic GAs (Figure S1). Interestingly, 3 of these amino acid insertions are outside the conserved S1-S5 regions, while two of the inserts are inside the S3 and S5 regions. The results suggest that the presence of these amino acid insertions may be an adaptation of SdGA to catalyze the reaction at low and medium temperatures.

## 4. Discussion

Numerous works have been carried out on the characterization of the structure and function of GAs from fungi and yeasts, however, a few enzymes from prokaryotic organisms have been studied. The first bacterial GA cloned and expressed in *E. coli* cells was that of *Clostridium* sp. [46]. *Clostridium* GA has maximum activity at pH around 5 and an optimal temperature close to 70 ° C. Subsequently, other GAs from thermophilic bacteria were characterized, such as those from *S. solfataricus*, *T. thermosaccharolyticum* or *T. tengcongensis*, which showed optimal temperatures between 60-90 ° C and an optimal acid pH, around 5 [5, 13] [47]. Other GAs from archaea such as *T. acidophilum*, *Picrophilus torridus* and *Picrophilus oshimae*, which showed optimal temperatures of around 90°C and pH 2.0 [48] or *Metanococcus jannaschii*, with optimal temperature and pH of 80°C and 6.5, respectively were recently described [12]. These GAs of archaea such as fungal GAs have higher specificity for larger substrates, such as starch, amylopectin, and glycogen, than for smaller maltooligosaccharides [49] [48] [50].

The common characteristic among the enzymes mentioned above is the high optimum temperature, which is desirable in various industrial processes such as the production of glucose and syrups [51]. However, there are very few reports on the characterization of GAs that work at low or medium temperatures. The latter would be relevant for certain processes such as cold starch hydrolysis for the production of bioethanol, as well as its use in the detergent and food industries, for the production of fine chemicals and in bioremediation processes [52]. [17, 40].

*Saccharophagus degradans* is a marine bacterium that could be defined as facultative psychrophilic or cold-tolerant, since although its optimal growth temperature is 37°C, it can grow in the range of 5-40°C [22]. As previously mentioned, this microorganism is related to the group of marine bacteria that degrade different complex polysaccharides with high efficiency that can be used as the only source of carbon and energy.

The results presented in this work indicate that SdGA has the ability to degrade maltose with a catalytic efficiency comparable to other described enzymes, including those from thermophilic organisms [46] (see Table S1) even at temperatures of around 10-20°C. Sequence alignment analysis revealed that this protein has 5 conserved regions (S1-S5), also present in other GAs previously described [53]. The highest sequence identity was observed with marine GAs from *G. polysaccharolyticus* and *G. agarilyticus* (67% and 66% identity, respectively), However, SdGA also shows a high degree of identity with GAs of other organisms that have the ability to grow at low temperatures such as many *Caulobacter* species (around 54-57% identity), or *Bevundimonas* species such as *B. mediterranea* (58% identity), *Pseudoalteromonas* species such as *P. arctica* (56% identity) and *Flavobacterium* sp. (56% identity) [41]. Furthermore, the alignment of the CD of SdGA with those from the GAs mentioned above showed an increase of around 6% of the identity (60 - 63% identity), which reinforces the fact that SdGA shares common characteristics with these enzymes.

The 3D model for SdGA shows a high structural conservation of the residues involved in the acarbose binding and active site. This protein was shown to be composed of a barrel-shaped CD (α/α)_6_ and an N-terminal β-sandwich domain similar to the thermophilic GA of *T. thermosaccharolyticum*, a representative member of the GH15 family. By homology modeling, we identified two glutamic acid residues in SdGA, E513 and E745, which are located at similar positions in other GAs, described as involved in the hydrolysis of substrates such as E438/E636 and E179/E400 from *T. thermosaccharolyticum* and *A. awamori* GAs, respectively [5, 54].

SdGA is active in an acidic environment, and this is characteristic of the catalytic mechanism of GAs. The widely accepted model for the action of GAs is the formation of oxocarbenium ions, which involves the transfer of protons to the glycosidic oxygen of the cleavable bond from a general acid catalyst, and a nucleophilic attack of water assisted by a general base catalyst [51, 55]. Unlike alpha-amylases where catalytic acid (proton donor) and base (nucleophile) are glutamate and aspartate residues, respectively, in GAs, glutamate residues act as acidic and basic catalysts [56, 57].

SdGA showed maximum activity at pH 6.0, similar to the optimal pH value of other microbial GAs, while the optimal temperature was 39°C [13, 47, 58–60]. Furthermore, the protein showed an activity greater than 50% in a wide range of pH (4.5 - 7) and temperatures (20-48°C), retaining around 30% of its activity at temperatures of 15°C, and a high catalytic efficiency, both desirable characteristics of cold-adapted enzymes.

All the kinetic parameters (*K*_m_, *k*_cat_ and *V*_max_) obtained for SdGA were within the previously reported values for microbial GAs (Table S1), however, SdGA showed no Ca^2+^ dependency and no inhibition at high concentrations of maltose. These data are comparable with those recently reported for the GA of *Caulobacter crescentus* [41]. This cold tolerant proteobacterium has a GA that shows an optimal temperature of 30°C and an optimal pH of 5.0, and is also active at low temperatures, showing a catalytic efficiency similar to the enzymes characterized from thermophilic organisms. The results obtained indicate that SdGA can be defined as a cold-adapted enzyme, with catalytic efficiencies comparable to other thermophilic GAs even at temperatures of 20°C.

Because most of the GAs that are currently being used in the starch saccharification stage have low activity at pH 6.0-6.5 [61], the use of SdGA that has optimal activity at pH 6.0 and it is active at low and medium temperatures, it represents a good alternative for this process because, for example, it would not be necessary to adjust the pH to 4.0-4.5, and thus, costs would be reduced. Furthermore, many GAs undergo substrate inhibition at high substrate concentrations [41, 62], while SdGA is not inhibited by maltose at concentrations greater than ten times the *S*_0.5_ value.

At temperatures above 45°C the SdGA activity decreased considerably. Thermal stability tests showed that SdGA is labile at temperatures above 45°C. These parameters are lower than those shown by fungal GAs, which have higher thermostability and show optimal temperatures between 40 - 70°C [63, 64]. Thermal stability analyses in the presence of metabolites showed that Ca^2+^ does not affect the stability of SdGA, while glycerol and acarbose partially stabilize the enzyme. In addition, Ca^2+^ also did not affect SdGA activity. The activity of fungal GAs is usually affected by Ca^2+^ [65, 66] and only a few GAs show activity in the absence of this cation [67]. From an industrial point of view, this is also a desirable feature since adding salts to any process would increase costs and / or could lead to unwanted side effects.

Finally, it was reported that at the structural level, cold-adapted enzymes are more flexible than their thermostable counterparts [43]. We found that SdGA shows similar flexibility than other GAs from psychrophilic or cold-tolerant organisms. Moreover, SdGA shows higher flexibility compared with thermophilic GAs. We also found that SdGA has a higher amino acid content respect to thermophilic GAs due to several insertions within its CD and also contains higher glycine content. It was reported that glycine residues provide flexibility to enzyme active sites [68]. These amino acid insertions, as well as the higher flexible residues and glycine content could explain not only the thermolability of SdGA, but also its ability to catalyze the reactions at low or mild temperatures. It was demonstrated that the features that make an enzyme more flexible, such as higher amino acid content, higher glycine content, and high flexible residue content near the active site compensates the low kinetic energy in cold environments [42, 43].

In summary, the novel GA from *S. degradans* has an overall structure like other GAs, showing a high structural conservation of the active site. SdGA is active over a broad range of pH and temperatures, it has no calcium dependency and shows no inhibition at high substrate concentrations, but an increased thermolability at moderate temperatures. We also found that SdGA is more flexible than its thermostable counterparts due to the higher content of flexible and glycine residues and a larger CD, due to several amino acid insertions in the vicinity of the conserved regions of this domain. These characteristics allow it to be classified as a cold-adapted enzyme. We propose that this novel SdGA, might have potential applications for use in different industrial processes that require an efficient alpha glucosidase activity at low/mild temperatures, such as biofuel production.

## Acknowledgements

NW (doctoral fellow), NH (postdoctoral fellow), MVB and DFGC (research scientists) are members of the National Research Council from Argentina (CONICET).

