## Supplemental files for "Characterization of SdGA, a cold-adapted glucoamylase from *Saccharophagus degradans*"

### Figure S1

```
                                                S1
AAK42610.1       242   -------NDYGEYDSILRRSLLILQS--HVQNNGAIVASLDT-----DIMKFNRDTYNYVWHRDAVFCILALE   300
WP_010900770.1   257   -----TPSLDSDTTALYRRSLFVTKS--HANDLGAIAASCDS-----DILKLSHDGYYYVWPRDASMAAYALS   317
AAC24003.1       304   ------NNFNGKANSLYYNSMMILKASEDKTNKGAYIASLSIPWGDGQR-DDNTGGYHLVWSRDLYHVANAFI   369
AAM25005.1       304   ------KNFGGEADELYYTSLMFLKASEDKTNKGAFIASLSIPWGEGQG-DENKGGYHLVWARDLYHIANAFI   369
WP_169043549.1   323   EPLNTMVKQTADDGKLLYTSAMVLKAQEDKTNAGALIASLSNPWGETVSAKEGSTGYKAVWVRDFYQVAMAFM   395
WP_149602793.1   325   KPLNTMVEQTADNGKLLYTSAMVLKAQEDKTHAGALIASLSNPWGETVSAKEGSTGYKAVWVRDFYQVAMAFM   397
WP_010557466.1   325   KPLNTMTEQTADNGKLLYTSAMVLKAQEDKTHAGALIASLSNPWGETVSAKEGSTGYKAVWVRDFYQVAMAFM   397
ABD79864.1       325   PNLPAMAAQATDGGKLAYTSAMVLKAQEDKSHAGALIASLSNPWGDIVPAVESSTGYKAVWPRDFYQVAMAFL   397
WP_047408988.1   296   TELPRLREASEDGGKLVQASALMLKVQEDRTHAGALIASLSNPWGDTVDATQSSTGYKAVWPRDFYQCAMALA   368
WP_154725846.1   315   TELPRIAEQATDGGKLAYASALMLKVQEDRTHAGALIASLSNPWGDTVDASKSSTGYKAVWPRDFYQVAMALA   387
WP_161638549.1   316   TELPRIAEQATDGGKLAYASALMLKVQEDRTHAGALIASLSNPWGDTVDASKSSTGYKAVWPRDFYQVAMALA   388

                                                                    S2
AAK42610.1       300   LMGYFDRSRQFFEFTRRLFT---------INGALFHKYTVDGHFGSTWHPWTLD---YLPIQEDETALVLYAL   361
WP_010900770.1   317   ISGHSETARRFFALMEDSLS---------EEGYLYHKYNVDGKIASSWLPHVMNGKSIYPIQEDETALVVWAL   381
AAC24003.1       370   AAGDVDSANRSLDYLAKVVK---------DNGMIPQNTWISGKP------------YWTGIQLDEQADPIILS   421
AAM25005.1       370   AAKDIDSANRALDFLAMVVE---------KNGFMPQNTWINGDP------------YWNGIQMDEQADPIILA   421
WP_169043549.1   396   ALGDTATAKTAFEYLEKVQVTSKTPGNNGDTGWFLQKTHVDGEL------------EWVGVQLDQTAMPIMLA   456
WP_149602793.1   398   ALGDTATAKTAFEYLEKVQVNSKTPGNKGDTGWFLQKTHVDGEL------------EWVGVQLDQTAMPIMLA   458
WP_010557466.1   398   ALGDTATAKTAFEYLEKVQVNSSTPGNNGDTGWFLQKTHVDGEL------------EWVGVQLDQTAMPIMLA   458
ABD79864.1       398   ALGDNETPKVAFEYLKKVQAGPDIEGYEGAPGWFLQKTHVDGTL------------EWYAVQLDQTAMPIMLG   458
WP_047408988.1   369   ALGDKQTPLAAFRYLPTVQVGAKTLGNKGDGGWFLQKSHVDGTP------------EWVGVQLDQTAMPIMLG   429
WP_154725846.1   388   ALGDRETPLAAFNYLPQVQVGPDTPGNTGVGGWFLQKTHVDGEL------------EWVAVQLDQTAMPIMLG   448
WP_161638549.1   389   ALGDKETPLAAFNYLPQVQVGPNTPGNTGAGGWFLQKTHVDGEL------------EWVAVQLDQTAMPIMLG   449
```

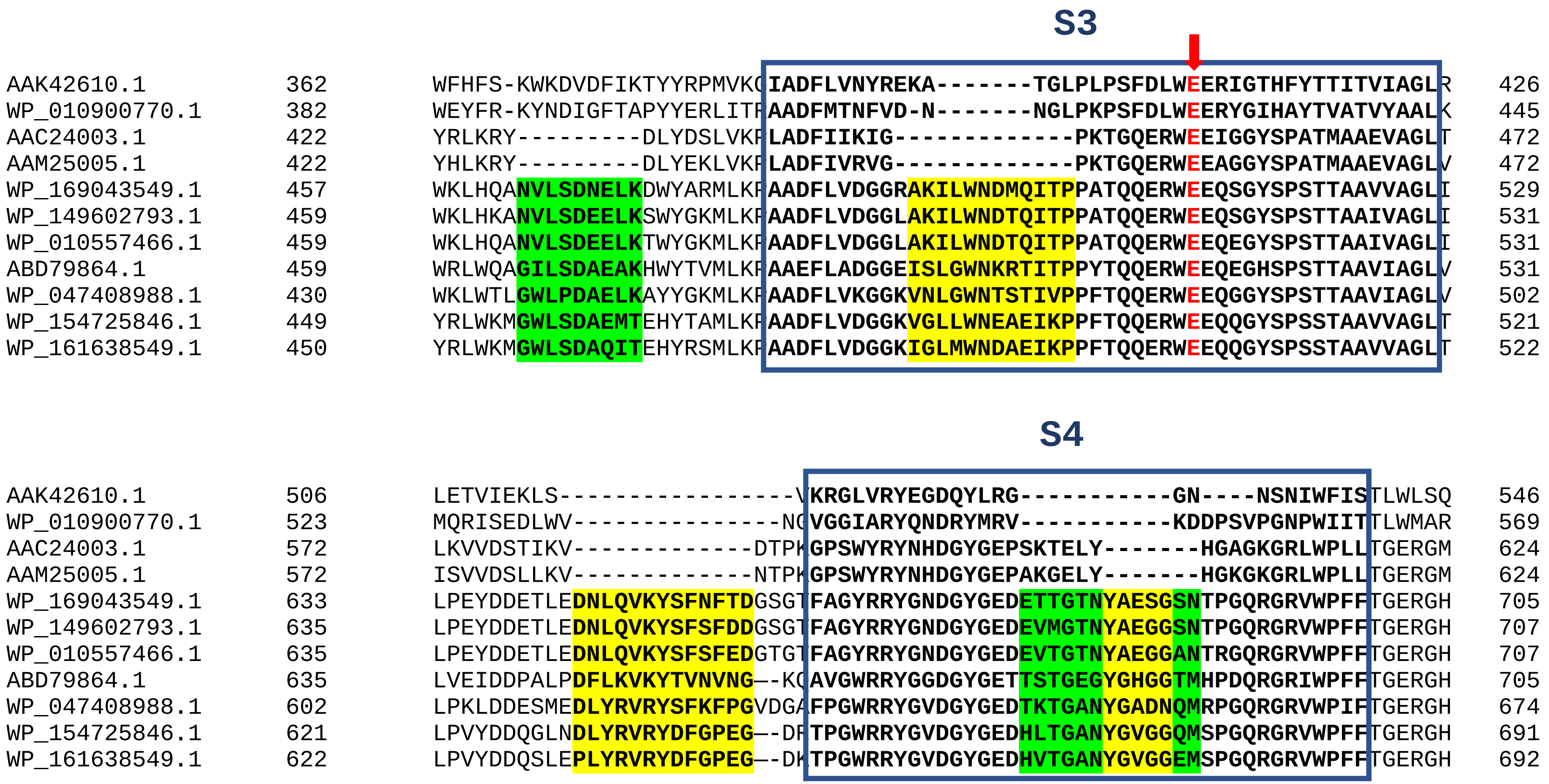
S3
AAK42610.1 362 WFHFS-KWKDVDFIKTYYRPMVKGIADFLVNYREKA-------TGLPLPSFDLWEERIGTHFYTTITVIAGLR 426
WP_010900770.1 382 WEYFR-KYNDIGFTAPYYERLITRAADFMTNFVD-N-------NGLPKPSFDLWEERYGIHAYTVATVYAALK 445
AAC24003.1 422 YRLKRY---------DLYDSLVKPLADFIIKIG-------------PKTGQERWEEIGGYSPATMAAEVAGLT 472
AAM25005.1 422 YHLKRY---------DLYEKLVKPLADFIVRVG-------------PKTGQERWEEAGGYSPATMAAEVAGLV 472
WP_169043549.1 457 WKLHQANVLSDNELKDWYARMLKPAADFLVDGGRAKILWNDMQITPPATQQERWEEQSGYSPSTTAAVVAGLI 529
WP_149602793.1 459 WKLHKANVLSDEELKSWYGKMLKPAADFLVDGGLAKILWNDTQITPPATQQERWEEQSGYSPSTTAAIVAGLI 531
WP_010557466.1 459 WKLHQANVLSDEELKTWYGKMLKPAADFLVDGGLAKILWNDTQITPPATQQERWEEQEGYSPSTTAAIVAGLI 531
ABD79864.1 459 WRLWQAGILSDAEAKHWYTVMLKPAAEFLADGGEISLGWNKRTITPPYTQQERWEEQEGHSPSTTAAVIAGLV 531
WP_047408988.1 430 WKLWTLGWLPDAELKAYYGKMLKPAADFLVKGGKVNLGWNTSTIVPPFTQQERWEEQGGYSPSTTAAVIAGLV 502
WP_154725846.1 449 YRLWKMGWLSDAEMTEHYTAMLKPAADFLVDGGKVGLLWNEAEIKPPFTQQERWEEQQGYSPSSTAAVVAGLT 521
WP_161638549.1 450 YRLWKMGWLSDAQITEHYRSMLKPAADFLVDGGKIGLMWNDAEIKPPFTQQERWEEQQGYSPSSTAAVVAGLT 522
S4
AAK42610.1 506 LETVIEKLS-----------------VKRGLVRYEGDQYLRG-----------GN----NSNIWFISTLWLSQ 546
WP_010900770.1 523 MQRISEDLWV---------------NGVGGIARYQNDRYMRV-----------KDDPSVPGNPWIITTLWMAR 569
AAC24003.1 572 LKVVDSTIKV-------------DTPKGPSWYRYNHDGYGEPSKTELY-------HGAGKGRLWPLLTGERGM 624
AAM25005.1 572 ISVVDSLLKV-------------NTPKGPSWYRYNHDGYGEPAKGELY-------HGKGKGRLWPLLTGERGM 624
WP_169043549.1 633 LPEYDDETLEDNLQVKYSFNFTDGSGTFAGYRRYGNDGYGEDETTGTNYAESGSNTPGQRGRVWPFFTGERGH 705
WP_149602793.1 635 LPEYDDETLEDNLQVKYSFSFDDGSGTFAGYRRYGNDGYGEDEVMGTNYAEGGSNTPGQRGRVWPFFTGERGH 707
WP_010557466.1 635 LPEYDDETLEDNLQVKYSFSFEDGTGTFAGYRRYGNDGYGEDEVTGTNYAEGGANTRGQRGRVWPFFTGERGH 707
ABD79864.1 635 LVEIDDPALPDFLKVKYTVNVNG--KQAVGWRRYGGDGYGETTSTGEGYGHGGTMHPDQRGRIWPFFTGERGH 705
WP_047408988.1 602 LPKLDDESMEDLYRVRYSFKFPGVDGAFPGWRRYGVDGYGEDTKTGANYGADNQMRPGQRGRVWPIFTGERGH 674
WP_154725846.1 621 LPVYDDQGLNDLYRVRYDFGPEG--DRTPGWRRYGVDGYGEDHLTGANYGVGGQMSPGQRGRVWPFFTGERGH 691
WP_161638549.1 622 LPVYDDQSLEPLYRVRYDFGPEG--DKTPGWRRYGVDGYGEDHVTGANYGVGGEMSPGQRGRVWPFFTGERGH 692

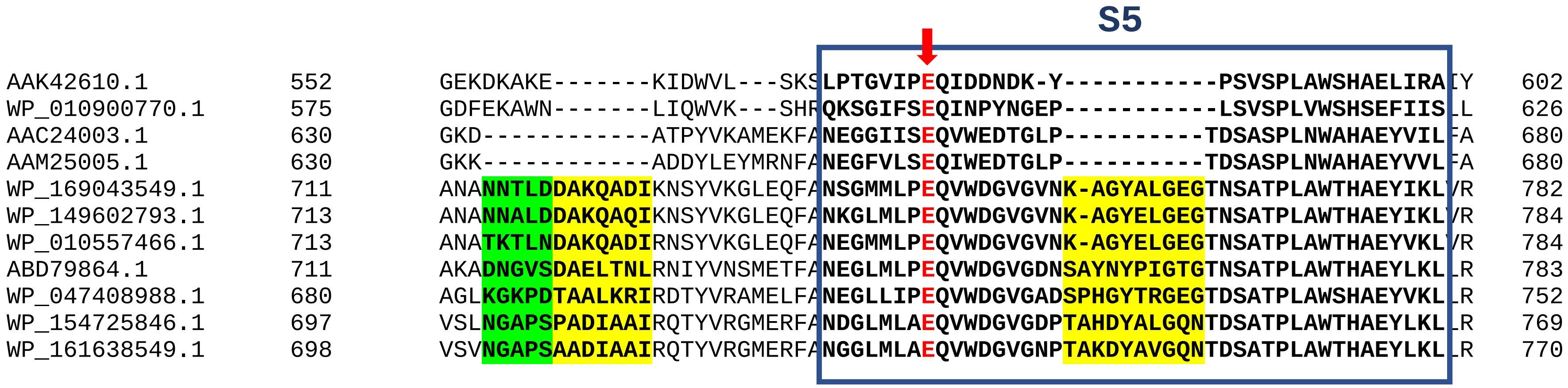


**Figure S1.** Sequence alignment of the conserved regions in the CD of SdGA with other GAs: AAK42610.1 *Sulfolobus solfataricus*; WP_010900770.1 *Thermoplasma acidophilum*; AAC24003.1 *Thermoanaerobacterium thermosaccharolyticum*; AAM25005.1 *Caldanaerobacter subterraneus* subsp. *tengcognensis*; WP_169043549.1 *Pseudoalteromonas arctica*; WP_149602793.1 *Pseudoalteromonas distincta*; WP_010557466.1 *Pseudoalteromonas marina*; ABD79864.1 *Saccharophagus degradans*; WP_047408988.1 *Caulobacter* sp; WP_154725846 *Brevundimonas mediterranea*; WP_161638549.1 *Flavobacterium* sp. S1-S5 are the conserved regions of the CD present in GAs. Glutamic acid residues involved in catalysis are marked with arrows. Amino acid insertions present in SdGA and pscicrophyllic and cold-adapted bacteria are shown in yellow. Insertions partially conserved in GAs are shown in green.

**Table S1. Comparison of kinetic parameters of various GAs with maltose as substrate**

| **Enzymes** | ***K*m (mM)** | ***k*cat ($s^{-1}$)** | **Specific activity (U/mg)** | **Temp. (°C)** | **pH** | **Reference** |
|---|---|---|---|---|---|---|
| SdGA | 11.5 | 8.9 | 6.1 | 39 | 6 | this work |
| *Picrophilus torridus* | n.a. | n.a. | 1.4 | 90 | 2 | (Serour and Antranikian 2002) |
| *Picrophilus oshimae* | n.a. | n.a. | 0.6 | 90 | 2 | (Serour and Antranikian 2002) |
| *Thermoplasma acidophilum* | n.a. | n.a. | n.a. | 75 | 5 | (Dock et al. 2008) |
| *Sulfolobus solfataricus* | n.a . | n.a. | 56.3 | 90 | 5.5 | (Kim et al. 2004) [3] |
| *Clostridium sp.* | 3.67 | 10.4 | n.a. | 25 | 4.5 | (Ohnishi et al. 1992) |
| *Rhizopus oryzae* | 0.53 | 9.3 | n.a. | 40 | 5.5 | (Mertens et al. 2010) |
| *Aspergillus awamori* | 0.72 | 8.8 | 20.6 | 50 | 4.4 | (Liu et al. 2000) |
| *Thermoactinomyces vulgaris R-47* | 0.1 | 16.9 | n.a. | 40 | 6.5 | (Ichikawa et al. 2004) |
| *Caldanaerobacter subterraneus subsp. tengcongensis MB4* | 13.4 | 149 | 80 | 75 | 5 | (Zheng et al. 2010) |
| *Caulobacter crescentus CB15* | 0.87 | 38.8 | n.a. | 30 | 5 | (Sakaguchi et al. 2014) |
| *Tetracladium sp.* | n.a. | n.a. | n.a. | 30 | 6 | (Carrasco et al. 2017) |

Ref: n.a.: not available

#### References Table S1


Carrasco M, Alcaino J, Cifuentes V, Baeza M (2017) Purification and characterization of a novel cold adapted fungal glucoamylase. Microb. Cell Factories 16(1):75 doi:10.1186/s12934-017-0693-x

Dock C, Hess M, Antranikian G (2008) A thermoactive glucoamylase with biotechnological relevance from the thermoacidophilic Euryarchaeon *Thermoplasma acidophilum*. Appl. Microbiol. Biotechnol. 78(1):105-14 doi:10.1007/s00253-007-1293-1

Ichikawa K, Tonozuka T, Uotsu-Tomita R, Akeboshi H, Nishikawa A, Sakano Y (2004) Purification, characterization, and subsite affinities of *Thermoactinomyces vulgaris* R-47 maltooligosaccharide-metabolizing enzyme homologous to glucoamylases. Biosci. Biotechnol. Biochem. 68(2):413-20 doi:10.1271/bbb.68.413

Kim MS, Park JT, Kim YW, Lee HS, Nyawira R, Shin HS, Park CS, Yoo SH, Kim YR, Moon TW, Park KH (2004) Properties of a novel thermostable glucoamylase from the hyperthermophilic archaeon *Sulfolobus solfataricus* in relation to starch processing. Appl. Env. Microbiol. 70(7):3933-40 doi:10.1128/AEM.70.7.3933-3940.2004

Liu HL, Doleyres Y, Coutinho PM, Ford C, Reilly PJ (2000) Replacement and deletion mutations in the catalytic domain and belt region of *Aspergillus awamori* glucoamylase to enhance thermostability. Protein Eng. 13(9):655-9

Mertens JA, Braker JD, Jordan DB (2010) Catalytic properties of two *Rhizopus oryzae* 99-880 glucoamylase enzymes without starch binding domains expressed in *Pichia pastoris*. Appl. Biochem. Biotechnol. 162(8):2197-213 doi:10.1007/s12010-010-8994-0

Ohnishi H, Kitamura H, Minowa T, Sakai H, Ohta T (1992) Molecular cloning of a glucoamylase gene from a thermophilic *Clostridium* and kinetics of the cloned enzyme. Eur. J. Biochem. 207(2):413-8 doi:10.1111/j.1432-1033.1992.tb17064.x

Sakaguchi M, Matsushima Y, Nankumo T, Seino J, Miyakawa S, Honda S, Sugahara Y, Oyama F, Kawakita M (2014) Glucoamylase of *Caulobacter crescentus* CB15: cloning and expression in *Escherichia coli* and functional identification. AMB Express 4(1):5 doi:10.1186/2191-0855-4-5

Serour E, Antranikian G (2002) Novel thermoactive glucoamylases from the thermoacidophilic Archaea *Thermoplasma acidophilum*, *Picrophilus torridus* and *Picrophilus oshimae*. Antonie van Leeuwenhoek 81:73-83

Zheng Y, Xue Y, Zhang Y, Zhou C, Schwaneberg U, Ma Y (2010) Cloning, expression, and characterization of a thermostable glucoamylase from *Thermoanaerobacter tengcongensis* MB4. Appl. Microbiol. Biotechnol. 87(1):225-33 doi:10.1007/s00253-010-2439-0
